# The number sense is an emergent property of a deep convolutional neural network trained for object recognition

**DOI:** 10.1101/609347

**Authors:** Nicholas K. DeWind

**Affiliations:** Department of Psychology, University of Pennsylvania, Philadelphia, PA

## Abstract

Humans and many non-human animals have the “number sense,” an ability to estimate the number of items in a set without counting. This innate sense of number is hypothesized to provide a foundation for more complex numerical and mathematical concepts. Here I investigated whether we also share the number sense with a deep convolutional neural network (DCNN) trained for object recognition. These *in silico* networks have revolutionized machine learning over the last seven years, allowing computers to reach human-level performance on object recognition tasks for the first time. Their architecture is based on the structure of mammalian visual cortex, and after they are trained, they provide a highly predictive model of responses in primate visual cortex, suggesting deep homologies. I found that the DCNN demonstrates three key hallmarks of the number sense: numerosity-selective units (analogous to biological neurons), the behavioral ratio effect, and ordinality over representational space. Because the DCNN was not trained to enumerate, I conclude that the number sense is an emergent property of the network, the result of some combination of the network architecture and the constraint to develop the complex representational structure necessary for object recognition. By analogy I conclude that the number sense in animals was not necessarily the result of direct selective pressure to enumerate but might have “come for free” with the evolution of a complex visual system that evolved to identify objects and scenes in the real world.

## Introduction

There is strong evidence for the nonverbal capacity to approximately enumerate the number of items in a visual array in adults, infants, and nonhuman animals (Merritt, DeWind, & Brannon, 2012). One hypothesis is that the number sense evolved to represent number because it was advantageous for a multitude of social and ecological functions (Dehaene, 2001). Several researchers have speculated that the number sense may have evolved to improve foraging efficiency (e.g., Barth, Kanwisher, & Spelke, 2003; Gallistel & Gelman, 2000; Piazza & Izard, 2009). For example, it is advantageous to be able to assess which tree has more ripe fruits before expending the energy to climb. Another hypothesis is that enumeration evolved for facilitating social interactions. For example, when deciding when to engage or back down in an altercation between groups, it would be advantageous to know how large your group is compared to your opponents’ group (McComb, Packer, & Pusey, 1994). Alternatively, individuals from flocking, herding, or schooling species may gain a reproductive advantage by joining larger groups of conspecifics (Agrillo, Dadda, Serena, & Bisazza, 2008).

Evolutionary hypotheses for the origins of psychological traits are notoriously difficult to test. Here I propose a novel source of evidence from *in silico* experiments examining artificial neural networks. Neural networks are a class of machine learning algorithms that are inspired by the structure of the nervous system and have the benefit of biological plausibility. Unlike a real brain, the factors shaping their representations of stimuli are known and under experimental control. So, if a particular representational structure, such as the number sense, can be observed in a neural network that has been optimized for other purposes, such as object recognition, we can conclude that the representational structure does not require direct natural selection to emerge.

Critical for this inference is the biological relevance of the neural network architecture, and neural networks have several commonalities with real brains. Most importantly, neural networks have units analogous to biological neurons, which have activation levels analogous to spike rates. The activation of one unit affects the activation of another via weighted connections analogous to a biological synapse. The networks are usually initialized with random synaptic weights, which are refined via a training process. The training process does not have a clear biological analogue, because biological systems do not start with randomly weighted synapses. Thus, the training combines the action of evolutionary refinement with learning over development to produce a complete “adult” network. Deep convolutional neural networks (DCNNs) are even more biologically relevant, incorporating two other aspects of mammalian visual cortex: iterative retinotopy and convolution. Retinotopy is preserved across several layers of the DCNN analogous to the iteration of the retinotopic map across V1, V2, and higher layers of mammalian visual cortex. Convolution mirrors the finding that similar feature detectors (e.g., an oriented Gabor wavelet filter) are found with different local receptive fields effectively tiling the retinotopic map within a layer. Although the inputs, architecture, and training algorithm of these networks are known, the activations of the units in response to novel stimuli are an emergent property requiring empirical investigation.

In addition to the noted similarities in architecture and function, there is also significant empirical evidence that the representational structure of DCNNs is very similar to the representational structure of the primate brain (Kriegeskorte, 2015). DCNNs explain significant variance in the response of neurons in macaque inferior temporal (IT) cortex (Yamins et al., 2014). Remarkably, DCNNs explain more variance in IT than any previously proposed model and approach the noise limit of how much variance can be explained in response to images of objects (Cadieu et al., 2014). There is also an interesting homology between the hierarchy of retinotopic maps in DCNNs and in mammalian visual cortex. The earlier layers of DCNNs have greater representational structure in common with early layers of visual cortex, and the later layers have more in common with higher levels of visual cortex (Khaligh-Razavi & Kriegeskorte, 2014). Although most studies of DCNN-brain homology focus on the similarity with IT, there is also evidence that the highest layers of DCNNs, those with the most abstract representations, also explain significant representational structure in the intraparietal and superior parietal regions in humans (Cichy, Khosla, Pantazis, Torralba, & Oliva, 2016). These parietal regions are similar to the areas that have been shown to encode numerosity in topographic maps across the cortical surface (Harvey, Klein, Petridou, & Dumoulin, 2013).

I identified three hallmarks of the number sense that could potentially be identified in the DCNN. The first is numerosity-selective neurons, which have been recorded in the dorsal-prefrontal cortex and the intraparietal sulcus (IPS) of monkeys (Nieder & Miller, 2004; Roitman, Brannon, & Platt, 2007; Viswanathan & Nieder, 2013) and in the medial temporal area in humans (Kutter, Bostroem, Elger, Mormann, & Nieder, 2018). Indirect evidence of numerosity-selective neurons in human IPS has been found using functional MRI techniques (Eger et al., 2009; Harvey et al., 2013; Piazza, Izard, Pinel, Le Bihan, & Dehaene, 2004). Numerosity-selective neurons have the unique property of modulating their activity depending on the number of objects observed by the subject but are relatively invariant to changes in non-numerical visual features such as the size of the items or their location within the array. These neurons are thought to form the neurobiological instantiation of the number sense (Nieder, 2016).

A second hallmark of the number sense is ratio dependence: the difficulty in comparing the numerosity of two sets of objects is related to the ratio of the number of items in each (Cantlon & Brannon, 2006). Furthermore, the shape of the ratio effect is well described by a cumulative log-normal distribution with a single free parameter for the slope, *w* (Piazza et al., 2010, 2004).

A third hallmark of the number sense is ordinality. We know that five is not four and five is not six, and we also know that five is *between* four and six. Animal experiments have demonstrated that monkeys trained to choose the larger numerosity on sets with 1-4 items can generalize to ordering sets with 5-9 items (Brannon & Terrace, 1998). This generalization also extends to larger values (Cantlon & Brannon, 2006), and monkeys do not readily learn non-sequential orders of numerosities (Brannon & Terrace, 2000).

Here I explored the representations of a DCNN that was previously trained to categorize photographs of objects (Krizhevsky, Sutskever, & Hinton, 2012). I showed the network thousands of dot array images and recorded the responses of the virtual neurons or units, testing for the three hallmarks of the number sense. I reasoned that if these hallmarks were present in an artificial neural network that had no optimization to represent numerosity for its own sake, we might infer that evolution of the biological number sense may be an emergent property and not the result of direct selection.

## Results

The Krizhevsky et al. (2012) DCNN consists of five convolutional layers followed by three fully connected layers. I recorded the responses in each layer to 5,590 images of between 4 and 64 white dots on a black field (Figure 1). The stimuli evenly sampled a three dimensional stimulus space developed previously (DeWind, Adams, Platt, & Brannon, 2015), with systematic variation of the size and spacing of the array elements controlling for multiple non-numerical visual features. I tested for the effect of the number, size, and spacing of the array elements on the activation of all units in the network. Previous studies recording from neurons have used a p-value threshold to define numerosity-selective units (e.g., Nieder & Miller, 2004; Roitman et al., 2007; Viswanathan & Nieder, 2013). However, given the large number of samples and lack of experimental noise in the simulation, the p-value threshold was too sensitive (Figure S1); I therefore followed Stoianov & Zorzi (2012) by defining a numerosity-selective unit as one for which numerosity explained at least 10% of the variance in the validation data set, and size and spacing did not explain more than 1% variance. Magnitude-sensitive units were defined as those for which the combination of numerosity, size, and spacing explained at least 10% of variance. Units that fell into neither group were classified as non-magnitude-sensitive. For brevity, I have plotted layer-level analysis results for Layers 1, 4, and 7 to give a representative look at how coding evolves through the hierarchy of the DCNN; summary statistics are given for all layers.

**Figure 1.**
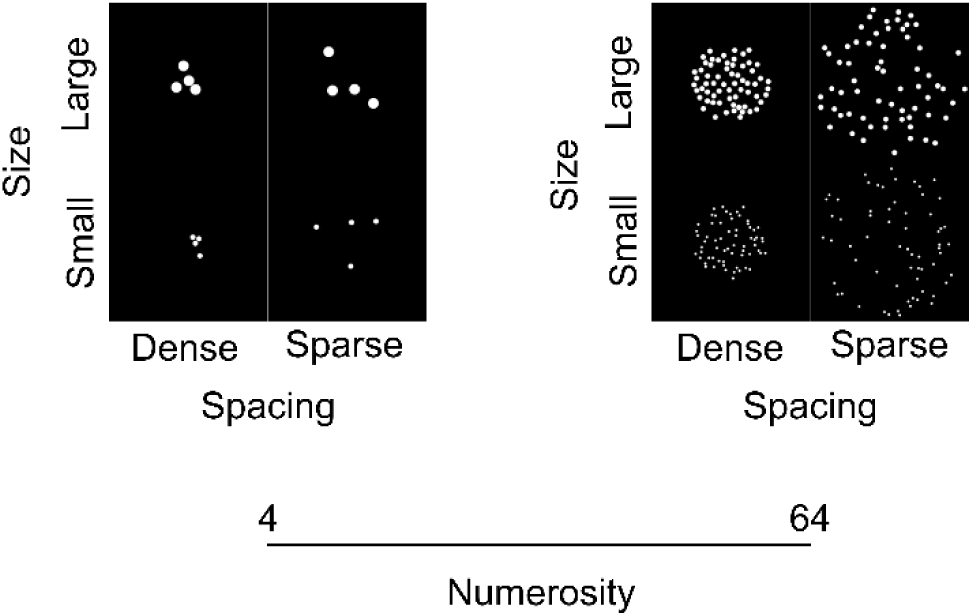
Example stimuli. The stimuli spanned a three-dimensional stimulus space varying the number, size, and spacing of array elements orthogonally. The smallest and largest size (bottom and top), the densest and sparsest spacing (left and right within each panel), and fewest and greatest number of items (left and right panel) of the primary stimulus set are shown.

Very little variance was explained by numerosity, size, or spacing in Layer 1, but in each subsequent convolutional layer more variance was explained (Figure 2A, first row). Variance explained peaked and plateaued in the three fully connected layers with an average of 41% variance explained by numerosity and an additional 22% explained by element size and spacing in Layer 7 (Figure 2B, top). In the fully connected layers almost every unit was sensitive to some combination of numerosity, size, and spacing (Figure 2A, first row; Figure 2B, middle). Apart from Layer 1, which only had two numerosity-selective units, the other layers consisted of 1.7% - 4.1% numerosity-selective units (Figure 2B, bottom), a prevalence that is on the same order of magnitude as some previous neural findings in monkeys (Viswanathan & Nieder, 2013), although is lower than other reports (Nieder & Miller, 2004; Roitman et al., 2007). This may reflect a true difference between the DCNN and the mammalian brain, or it may reflect overestimation in monkey studies as a result of failure to detect small effects of non-numerical visual features on inherently noisy neuronal firing rates combined with a reliance on p-values for neuronal classification. Fully connected layers averaged slightly higher prevalence of numerosity-selective units than the convolutional layers, but there was no clear pattern of increasing prevalence after Layer 1. For example, Layer 4 and Layer 7 both consisted of 2.6% numerosity-selective units. Thus, although the variance explained by numerosity rose dramatically across the layers, the variance explained by size and spacing also rose, and the proportion of numerosity-selective units was relatively flat. Figure S2 plots the activations of example numerosity-selective units.

**Figure 2.**
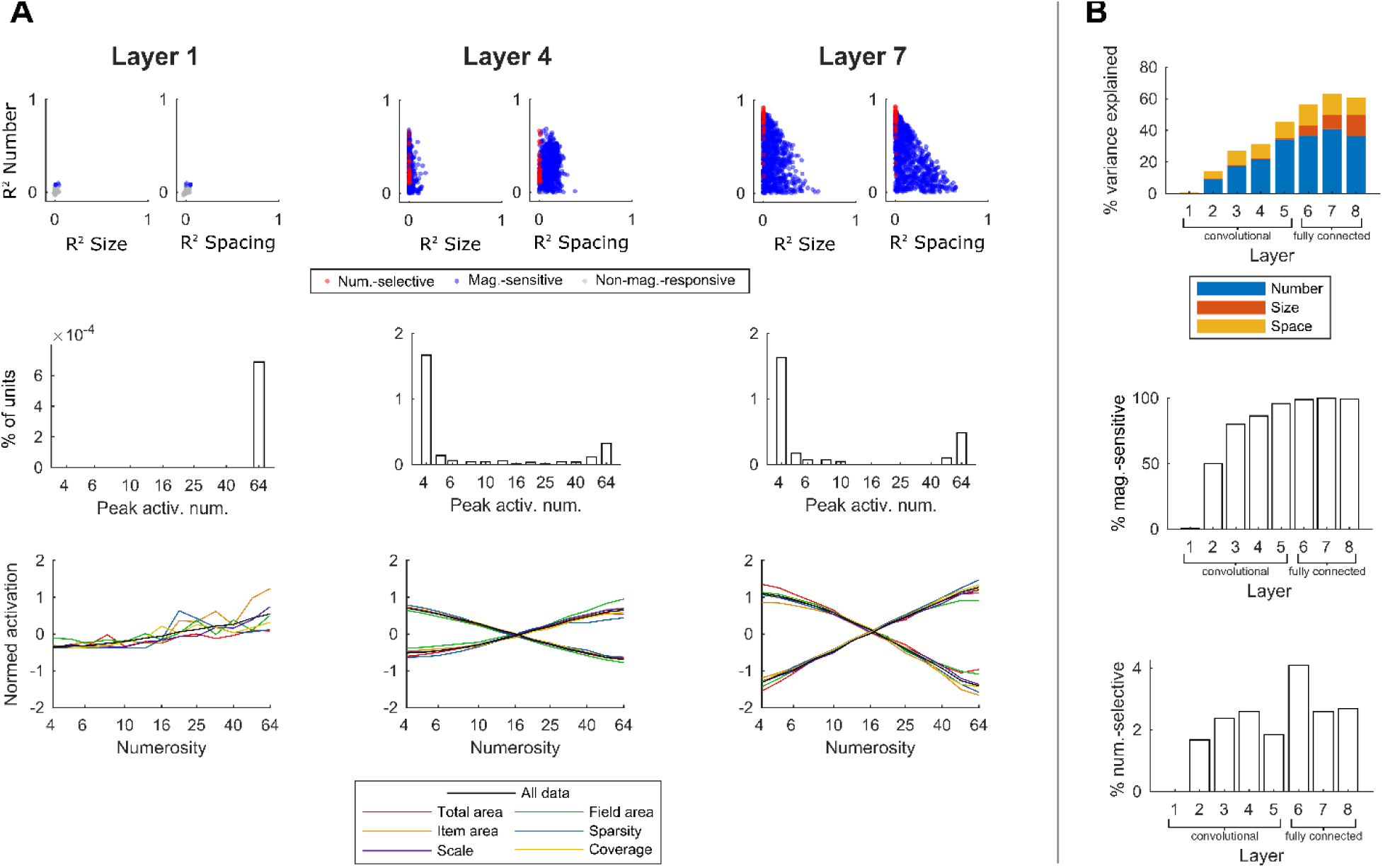
Numerosity explains a large portion of variance in DCNN responses, and numerosity-selective units exist in all layers. (A) Layer specific analyses for Layers 1, 4, and 7 (columns). The first row plots the predictive accuracy of the number model plotted against the predictive accuracy of the size and of the spacing models for 1,000 randomly selected units from each layer. The color of the units denotes how they were classified based on variance explained. The second row is a histogram of the numerosities that elicited maximum activation from the numerosity-selective neurons. The third row plots the average response of all the numerosity-selective units in each layer that were monotonically increasing and monotonically decreasing (in Layer 1 there were only two numerosity selective units and both were monotonically increasing). The black lines are the average across all images, and the colored lines are averages within the subsets of images in which numerosity varied while a non-numerical visual feature was held constant. These visual features were total area (red), item area (orange), field area (green), sparsity (blue), scale (purple), and coverage (yellow). Activations were z-scored within unit before averaging. (B) Summary statistics for all layers. The top plot shows the mean variance explained in unit activation in each layer by the number, size and spacing models; the middle plot shows the proportion of each layer comprised of magnitude-sensitive units; the bottom plot shows the proportion comprised of numerosity-selective units.

Previous research has found numerosity-selective neurons of two types: tuned (Nieder & Miller, 2004) and monotonic (Roitman et al., 2007). Tuned neurons have a normal-distributed activation profile across numerosities with a peak at a preferred numerosity, whereas monotonic neurons steadily increase or decrease activity across numerosities. I plotted a histogram of the numerosity that caused peak mean activation in each numerosity-selective unit (Figure 2A, middle row). Most units responded maximally to either the largest or smallest numerosity displayed, consistent with monotonic coding, although some neurons responded maximally to intermediate numerosities consistent with tuned coding.

Many previous studies demonstrating tuned representations of neurons in monkeys were conducted using arrays with 1-5 items (e.g., Nieder & Miller, 2004). I generated a second stimulus set consisting of numerosities 1-5 to explore whether units that were maximally activated by four items in the primary stimulus set were actually tuned to smaller numerosities. Across both stimulus sets and all layers, 29% of numerosity-selective units were tuned and 71% were monotonic (responding maximally to 1 or 64; Figure S3). Above Layer 1, which had too few numerosity-neurons to be relevant, there was no discernable pattern by which the proportion of types changed across the DCNN hierarchy (Figure S3B).

Many non-numerical visual features typically vary with the number of items in a visual array. It is not possible to orthogonalize all of them to numerosity simultaneously in a single stimulus set; however, within subsets of the images, a particular non-numerical visual feature was fixed while numerosity varied over its entire range. To demonstrate that numerosity and not one of the non-numerical visual features was driving changes in activation of the numerosity-selective units, I plotted the mean activation within a layer to each stimulus control subset, as well as to the entire stimulus set (Figure 2A, bottom row). Regardless of which non-numerical feature was held constant, these units responded consistently to changes in numerosity, confirming that they were indeed numerosity-selective.

To examine the “behavior” of the DCNN number sense, a support vector machine (SVM) linear classifier was trained to estimate numerosity from the activations of all units in each layer. The classifier was trained to discriminate each pair of numerosities and tested on held-out validation data. Figure 3A (first row) shows the classifier accuracy on all pair-wise combinations of numerosity. Classifier performance goes from near-chance performance in Layer 1 to highly accurate in the fully connected layers. Figure 3A (second row) shows the same accuracy data averaged within pairs of similar numerical ratio. In animals and humans, the function relating the logarithm of the numerical ratio to the accuracy of discrimination closely follows a cumulative normal curve with a single free parameter, *w*, proportional to the reciprocal of the slope. The *w* term denotes the acuity of the number sense, with smaller *w* indicating better performance (Piazza et al., 2004). I fit this psychometric function to the classifier accuracy in each layer (Figure 3A, second row). The *w* failed to converge to a finite positive value in Layer 1, because there was a slight negative trend across numerical ratio. In the other convolutional layers the best fit *w* was very large by behavioral standards; however, in the fully connected layers the *w* jumped to human-like levels (Figure 3B, top). The behavioral model was a very poor fit to the accuracy data in the convolutional layers, because there was not a clearly defined monotonic effect of numerical ratio on classification performance. However, the model was an almost perfect fit in the fully connected layers (Figure 3B, bottom). The classifier performance was so good in the fully connected layers that most of the ratios tested were close to ceiling. To get a more precise estimate of *w* and to be sure that the high degree of model fit was not due to a simple ceiling effect, I calculated DCNN activations to another stimulus set with more difficult ratios that had been used in a human behavioral experiment (DeWind et al., 2015). The *w* term was in the range of human performance and the psychometric function fit almost perfectly (Figure S4).

**Figure 3.**
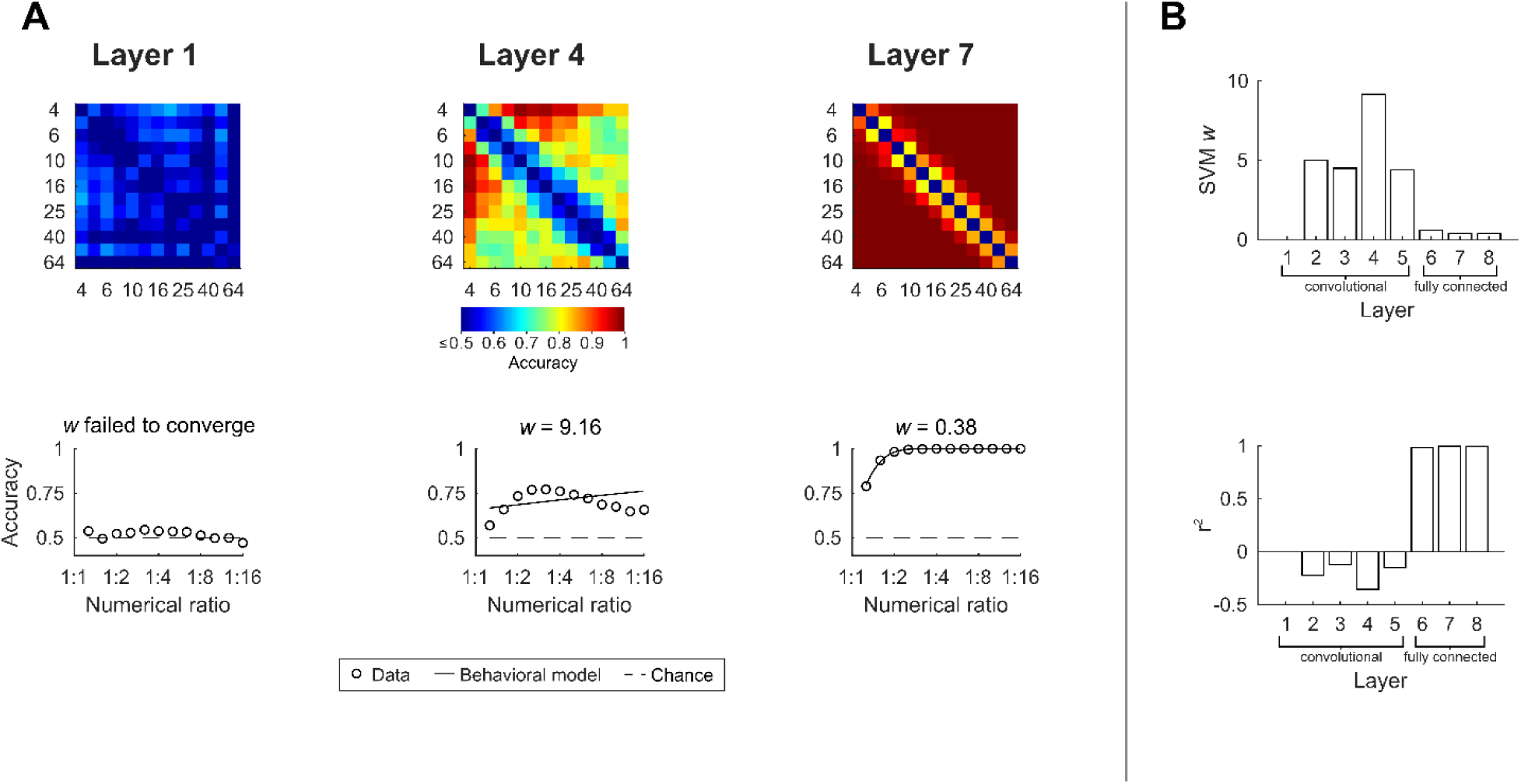
Human-like performance emerges through the DCNN hierarchy. (A) The first row plots heat-maps of SVM classifier accuracy for all pair-wise comparisons of numerosity. The second row shows the same data averaged over similar ratios. The line indicates the fit psychometric function, and *w* indicates the fit value of the single free parameter. (B) The top plot shows the *w* for each layer. The bottom plot shows the signed r^2^ of the fit of the psychometric function. The r^2^ can be negative if the residual sum of squared error is greater than the sum of squared error of the mean.

To ensure that the SVM classifier was discriminating on the basis of numerosity as opposed to another visual feature that was correlated with numerosity, the classifier was tested on subsets of the stimuli that controlled for different non-numerical visual features (Figure S5). In Layer 1 performance was close to chance across all subsets. In Layer 4 performance was above chance in every condition, but accuracy varied across conditions indicating that the classifier was using a combination of numerosity and other visual features, notably sparsity. In Layer 7 performance was very high and there was almost no difference in accuracy across conditions, indicating that the classifier was correctly categorizing images based on numerosity.

Previous work has expanded on the Piazza et al. (2004) psychometric function to allow for the simultaneous assessment of numerical acuity and the influences on non-numerical visual features in an ordinal comparison task (DeWind et al., 2015). In short, a multi-dimensional cumulative normal curve is fit to choice performance over the logarithm of the numerical ratio, the size ratio, and the spacing ratio. The coefficient estimates (β_Num_, β_Size_, and β_Spacing_) indicate which visual features influenced choices in an ordinal comparison task. I fit this model to the SVM pair-wise stimulus classifications in each layer and compared the results to 20 human participants’ choice data collected and reported previously (DeWind et al., 2015) (Figure S6). Layer 1 was largely unaffected by any stimulus feature and the other convolutional layers were influenced by numerosity and sparsity by approximately equal degree. The three fully connected layers performed unbiased numerosity comparisons (unaffected by the size or spacing of the elements), and Layers 7 and 8 were within the human distribution of performance.

The pair-wise classification of numerosities demonstrates that the DCNN represents differences in array element number. However, it is unclear if this representation is only categorical, in the same sense that the network might differentiate a dog from a cat, or if it is ordinal in the sense that all numerosities are represented along a single dimension in their proper order. To explore this question quantitatively, another SVM classifier was trained to classify numbers as greater or less than sixteen. The classifier was trained on numerosities 4, 6, 10, 25, 40, and 64, and the numerosities 5, 8, 13, 20, 32, and 51 were held out for testing. Classification performance on held-out data was above chance in all layers, demonstrating generalization to novel numerosities and some degree of ordinality (Figure S7).

Finally, I explored the ordinality qualitatively by plotting the first two principal components of the unit-by-image matrix of activations for each layer (Figure 4). Layer 1 lacks a discernible pattern across numerosities. Layer 4 has a clear progression, but the larger numerosities become more diffuse and curve through the space, making linear classification more difficult. By Layer 7 a relatively linear “mental number line” emerges.

**Figure 4.**
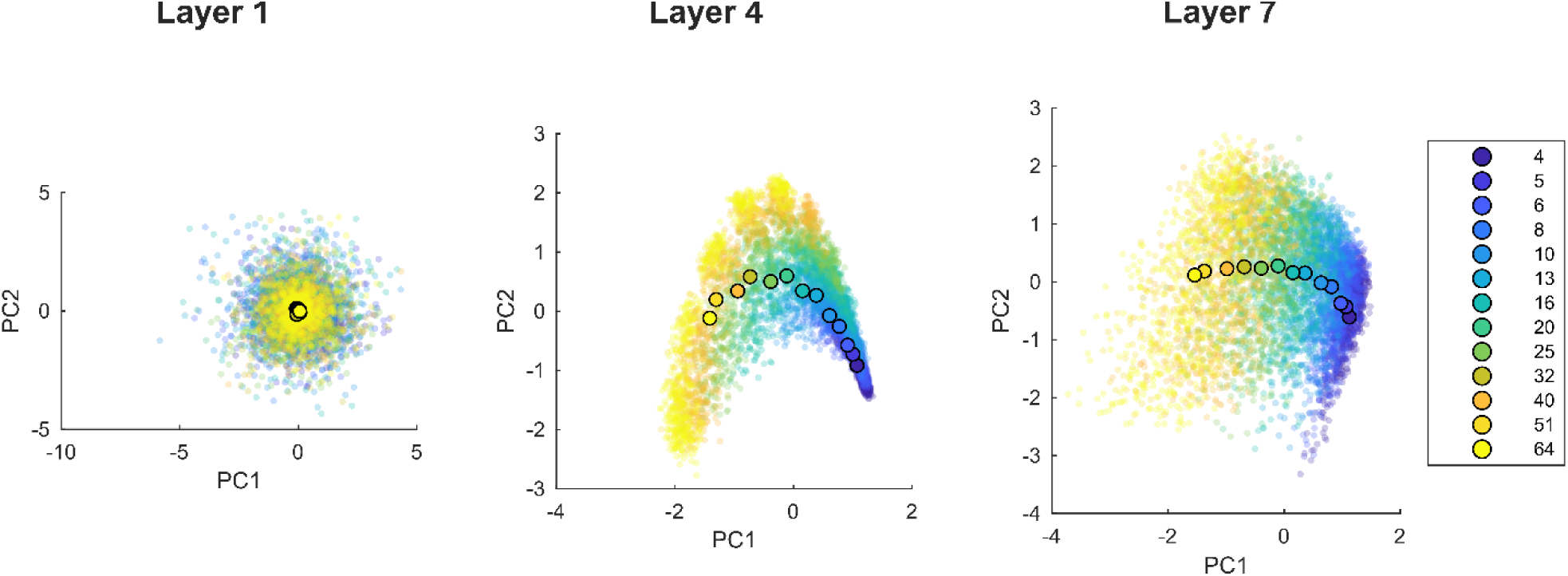
The “mental number line” emerges across the DCNN hierarchy. The first two principal component scores from the unit-by-image matrix of activations in response to all images in the primary stimulus set color coded by numerosity (n = 5,590). Numerosity means plotted in larger markers with black edges.

## Discussion

The DCNN tested here was trained for object recognition and was never explicitly optimized for enumeration (Krizhevsky et al., 2012). Nevertheless, I demonstrated that the network has an emergent sense of number remarkably similar to the biological number sense characterized in humans and non-human animals. The DCNN number sense must be the result of some combination of the network architecture and the constraint to develop the complex representational structure necessary for object recognition. The network architecture is inspired by the mammalian visual system, and empirical research has demonstrated deep homologies between the DCNN and primate visual cortex (Kriegeskorte, 2015). Thus, I conclude that the biological number sense may not have been directly optimized by natural selection to enumerate but rather “came for free” with the evolution of a complex visual system that evolved to identify objects and scenes in the real world.

A spandrel is an architectural term for the triangles created by the meeting of two arches or an arch and a wall. In their influential paper, Gould and Lewontin (1979) rhetorically ask the purpose of a spandrel and conclude that it has no purpose but rather is a byproduct of arches, which have the critical purpose of supporting walls and roofs over openings. By analogy, they argue that many phenotypes that have been hypothesized to be adaptations to the environment may be byproducts of some other adaptive pressure or the result of a constraint imposed by the overall structure or development of the organism (the “body plan” or “Bauplan”). The number sense may be an evolutionary spandrel: complex visual recognition constitutes the arch, and the number sense is a byproduct. However, an alternative is that the number sense constitutes a subsection of the arch itself; object recognition may be improved by virtue of numerosity-selective neurons, and enumeration – perhaps of sub-components of the objects themselves – is an important component of the algorithm for object recognition instantiated by the DCNN. Interventional experiments manipulating the activations of the DCNN may help clarify these possibilities. More generally, it is useful to consider both the adaptive and non-adaptive explanations of cognitive traits, and DCNNs can provide a novel avenue for exploring the evolutionary causes of neural representations.

In addition to inferences about evolution, the details of the DCNN findings have important implications for the refinement of computational models of the number sense. Previous models posit three hierarchically organized levels of processing: a normalization stage, a summation stage containing units that monotonically vary with numerosity, and a number-field stage containing units tuned to specific numerosities (Dehaene & Changeux, 1993; Verguts & Fias, 2004). Two predictions of these models are that numerosity-selectivity emerges late in cortical processing and that monotonic neurons precede tuned neurons in that processing. However, in the DCNN there are tuned and monotonic numerosity-selective units as early in the hierarchy as Layer 2. Layer 2 of the Krizhevsky et al. DCNN has the strongest representational similarity to early visual cortex of any layer (Khaligh-Razavi & Kriegeskorte, 2014; Kriegeskorte, 2015). This suggests numerosity-selectivity may develop earlier in the visual stream than previous models of the number sense predict and does not depend on earlier stages of cortical processing. This is consistent with the recent empirical finding that numerosity is encoded in early visual cortex (DeWind, Park, Woldorff, & Brannon, 2018; Eger, Pinel, Dehaene, & Kleinschmidt, 2015; Park, DeWind, Woldorff, & Brannon, 2016). Exactly how the well-characterized receptive field properties of neurons in early visual cortex might give rise to numerosity-selectivity requires further investigation.

Another novel insight from the DCNN is that numerosity-selective units – tuned or monotonic – were not sufficient to give rise to all the behavioral signatures of the number sense. Even though both the proportion of numerosity-selective units and the relative proportion of tuned to monotonic numerosity-selective units were relatively flat across the layers of the DCNN, there was a dramatic shift in population-level numerosity encoding between the convolutional and fully connected layers. This change was marked by a clearer ratio effect that extended across all ratios and numerosities, a higher overall classification accuracy very similar to human performance, and a shift from partial reliance on sparsity to total reliance on numerosity for classification.

If the proportion and type of numerosity-selective units are not sufficient to generate the behavioral signatures of the number sense, then what explains the emergence of the number sense in the later layers of the DCNN? One contributing factor is likely variance explained by numerosity. Although both the early and later layers of the DCNN had units that varied selectively with numerosity, units in the later layers varied much more with numerosity relative to their total variance. Another factor is probably related to full connectedness of Layers 6, 7, and 8. Convolutional layers retain retinotopic organization, and the receptive fields of the units are spatially restricted. As a result, their representation of numerosity at the unit level is necessarily local. The individual units in the fully connected layers receive input from the entire image. This might be critical for the computational steps necessary for the emergence of the number sense, including the ability to ignore non-numerical local features such as sparsity. This is consistent with the finding that numerosity can be decoded from early visual cortical layers, but classification fails to generalize to changes in the presentation location of the stimuli (Eger et al., 2015). Thus, a robust representation of numerosity sufficient for all the behavioral signatures of the number sense may only be possible in regions of cortex that are responsive to visual stimuli, but not retinotopically organized.

## Conclusion

Here I demonstrated that the number sense exists in a DCNN trained for object recognition. Because the network was not trained on enumeration, the number sense can be considered an emergent property of the network: the result of the optimization for object recognition and inherent architecture. The DCNN has many similarities to and empirically demonstrated homologies with the primate visual cortex, and so this finding serves as evidence that the biological number sense might similarly be an emergent property of a sufficiently robust representation of the world to guide behavior and of the cellular and physiological architecture of the brain. DCNNs offer a unique tool for testing evolutionary hypotheses of neural function and provide a uniquely transparent model system for testing computational theories of perception.

## Acknowledgments

I would like the acknowledge Kailen Mooney, Emily Szkudlarek, Stephanie Bugden, and Rosa Rugani for helpful discussions. I would also like to acknowledge Monica Carlson and Elizabeth Brannon for their feedback on drafts of the manuscript.

## Methods

### Experimental model

The DCNN was a pretrained network designed to perform object recognition (Krizhevsky et al., 2012). The network is divided into eight layers. The first five are convolutional and the last three are fully connected. The units in the convolutional layers can be classified by their location in three dimensions. The first two describe their x and y coordinates related to the pixel locations of the image (or previous layer). The third describes the filter function that is convolved with the image (or previous layer). The analyses presented differentiated between units of different layers but did not differentiate for the x and y locations or filter channel.

### Method details

#### Stimulus features and general characteristics of the stimulus space

There were three stimulus sets used in this manuscript, the 4-64 stimulus set, the 8-32 stimulus set, and the 1-5 stimulus set. Unless otherwise specified, the analyses described in the Results section were performed on the 4-64 stimulus set. All three stimulus sets sampled the three-dimensional stimulus space described previously (DeWind et al., 2015). Stimuli were generated using custom MATLAB scripts.

Several visual features of the stimuli are referred to in the manuscript; they are rigorously defined here. Item area refers to the area of each dot in the array. Total area is the item area multiplied by the number of items. Field area refers to the area of the invisible circle (the “field”) within which the items were drawn. Sparsity is the field area divided by the number of items. Coverage is total area divided by the field area. Scale is item area multiplied by field area. In each stimulus set, all of these features varied over the same number of octaves (doublings) as number. For example, in the 4-64 stimulus set the smallest and largest numerosities differed by 16×; total area, item area, field area, sparsity, coverage, and scale also varied by 16× between their smallest and largest values. As a result, the analyses were equally likely to detect a change in network activation due to change in any of these features, and the tests were not biased towards detecting an effect of numerosity.

To construct an array, the number of items, the item area, and the field area were specified. Items were randomly placed in the image such that they did not overlap with each other or the edge of the field.

Two other features, size and spacing, were important to the overall construction of the stimulus sets. These features are designed to account for the non-independence of the other features, which results in the problem that not all features can be simultaneously orthogonalized to number. For example, because total area is equal to number multiplied by item area, these features are not independent. Although there are three features of interest, there are only two degrees of freedom. Size represents the single degree of freedom remaining after number is fixed. Intuitively, if the number of items is set and one increases or decreases the size of the items in the array, then the total area and item area both vary together. Similarly, number, sparsity, and field area are not independent. Spacing represents the single degree of freedom remaining after number is fixed. Size is defined as the item area multiplied by the total area, and spacing is defined as the sparsity multiplied by the field area. These seemingly arbitrary definitions are designed so that when log number, log size, and log spacing are orthogonal and vary by the same factor (e.g., 16x), then item area, total area, sparsity and field area will all be equally but only partially correlated with number and vary by the same factor. Coverage and scale will also vary by the same factor and will be orthogonal to number. These attributes of all three stimulus sets make them ideal for maximizing the power to detect effects of each stimulus feature on activation without biasing the test toward finding an effect of a particular stimulus feature over another.

#### 4-64 stimulus set

Number, size and spacing were orthogonally varied over 13 levels covering four octaves (16x difference between smallest and largest values; Figure S8). The levels of numerosity were 4, 5, 6, 8, 10, 13, 16, 20, 25, 32, 40, 51, and 64. In order to reduce the number of stimuli, each level of number was sampled at every other level of size and spacing (and vice-versa). In the 3D stimulus space defined by number, size, and spacing, this resulted in two nested cubic grids of stimuli, one consisting of 7^3^ and one of 6^3^ stimuli, resulting in 559 unique locations. This sampling of the stimulus space is analogous to the way two nested rectangular grids of stars are arranged on the flag of the United States of America (6×5 + 5×4 = 50 stars). The levels of each feature are approximately evenly distributed along a log-scaled axis with slight deviation necessary for rounding. Ten images were generated at each stimulus space location resulting in 5,590 total images.

#### 8-32 stimulus set

This stimulus set was used previously for a human behavioral experiment (DeWind et al., 2015). This set is very similar to the 4-64 stimulus set in that each feature varied over 13 levels, but in the 8-32 set only two octaves were sampled. As a result, each level was more closely spaced on the log-scale feature axes, and the numerical ratios were closer. The numerosities sampled were 8, 9, 10, 11, 13, 14, 16, 18, 20, 23, 25, 29, and 32. There were 559 unique stimulus space locations sampled and ten images generated at each location resulting in 5,590 images.

#### 1-5 stimulus set

This stimulus set sampled the numerosities 1, 2, 3, 4, and 5. Size and spacing were also sampled at 5 levels at approximately equal distances on a log scale (not evenly spaced). In this stimulus set, every size and spacing level was sampled at every numerosity level (and vice-versa). This resulted in 5^3^ or 125 unique stimulus space locations. Ten images were generated at each location resulting in 1,250 images.

### Quantification and statistical analysis

#### Activations

Activations in response to the images were obtained using the Neural Network Toolbox (version 11) for MATLAB (version R2017b, Mathworks). The response of every unit to every image in the three stimulus sets was recorded.

#### Unit classification

For each unit I fit three regression models predicting activation, one each for numerosity, size, and spacing. Each regression model used regressors dummy coding for all 13 levels of the relevant feature. Models were fit to half the data and predictions were made for the held-out validation set. The correlation coefficient (Pearson’s r) between the vector of predicted activations and the vector of observed activations was calculated and squared with sign preserved. The p-values for these correlations are plotted in Figure S1. These values are plotted in Figure 2A (top row), and the means of these values in each layer are plotted in Figure 2B (top). Units were classified as numerosity-selective if the number model R^2^ was greater than 0.1 and the size model R^2^ and spacing model R^2^ were both less than 0.01. Units were classified as magnitude-sensitive if the sum of the three models’ R^2^ was greater than 0.1.

#### Peak activation numerosity

For each numerosity-selective unit, the peak activation numerosity was simply the numerosity that elicited the greatest mean activation (Figure 2A, middle row). In Figure S3A, units that had peak activation to 4 in the 4-64 stimulus set were re-classified as the peak activation the 1-5 stimulus set, unless the peak was at 5, in which case they were still classified as peaking at 4. Monotonic numerosity-selective units were defined as those with peak activations for 1 or 64. Tuned numerosity-selective units were defined as those with peak activations for any intermediate value (Figure S3B).

#### Classifiers

The dimensionality of the image-by-unit matrix of activations was reduced using PCA. The PCs accounting for 99% of the variance in activation were z-scored and used for classification. SVM classifiers (linear kernel) were trained on all pair-wise comparisons of numerosity using half the activation data (MATLAB Statistics and Machine Learning Toolbox version 2017b). The classifiers were then tested on the held-out validation activation data and accuracy was recorded.

For the classification of stimuli in the stimulus-control-subsets (Figure S5), these same classifiers were tested on the stimulus-control-subset portions of the validation data separately.

For the test of ordinality (Figure S7), a single classifier was trained to distinguish between stimuli with numerosity greater than 16 and less than 16 over portion of the training set with numerosities 4, 6, 10, 25, 40, and 64. It was tested on the subset of the validation data with numerosities 5, 8, 13, 20, 32, and 51.

#### Psychometric functions

We fit two psychometric functions to the pair-wise SVM classification data. The first was based on Piazza et al. (2004). A GLM with a single regressor (log_2_ of numerical ratio) was fit to the accuracy data using a binomial error distribution and a probit link function. The *w* term was defined based on regression coefficient as below:

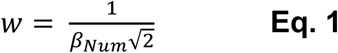

The second psychometric function was based on DeWind et al. (2015). A GLM was fit to the binary choice data with regressors for log_2_ of numerical ratio, log_2_ of size ratio, and log_2_ of spacing ratio. The error distribution was binomial and a probit link function was used.

