## Supplemental Figures for "The number sense is an emergent property of a deep convolutional neural network trained for object recognition"

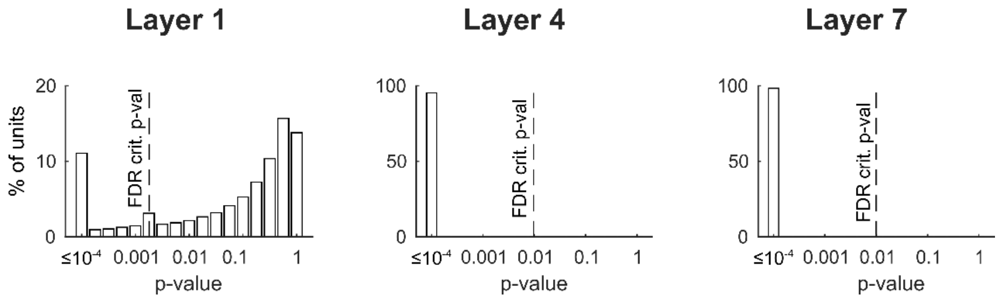


**Figure S1. Numerosity predicts unit activation to held out validation-set images in all layers.** A histogram of p-values for the correlation between activations predicted by the numerosity model and emperical activations. Dotted line indicates the FDR critical p-value for an expected false discovery rate of 1% (Benjamini & Hochberg, 1995).

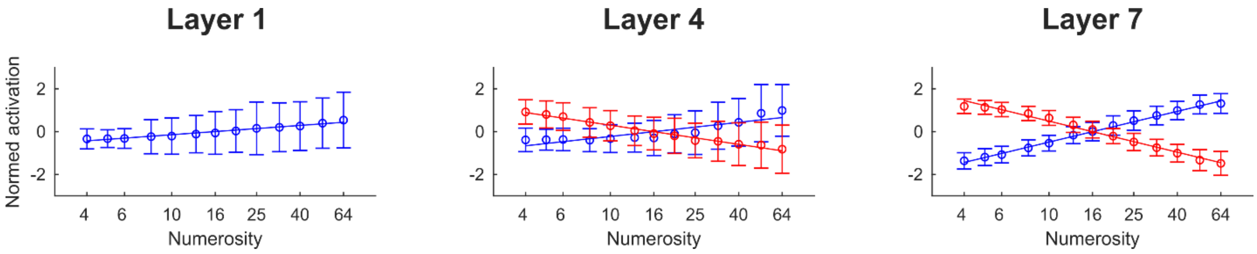


**Figure S2. Example numerosity-selective units.** Activation plotted across numerosity of one monotonic increasing numerosity-selective unit from Layer 1 and one monotonic increasing and one monotonic decreasing numerosity selective-units each from Layers 4 and 7. Activation was z-scored. Error bars indicate standard deviation over images.

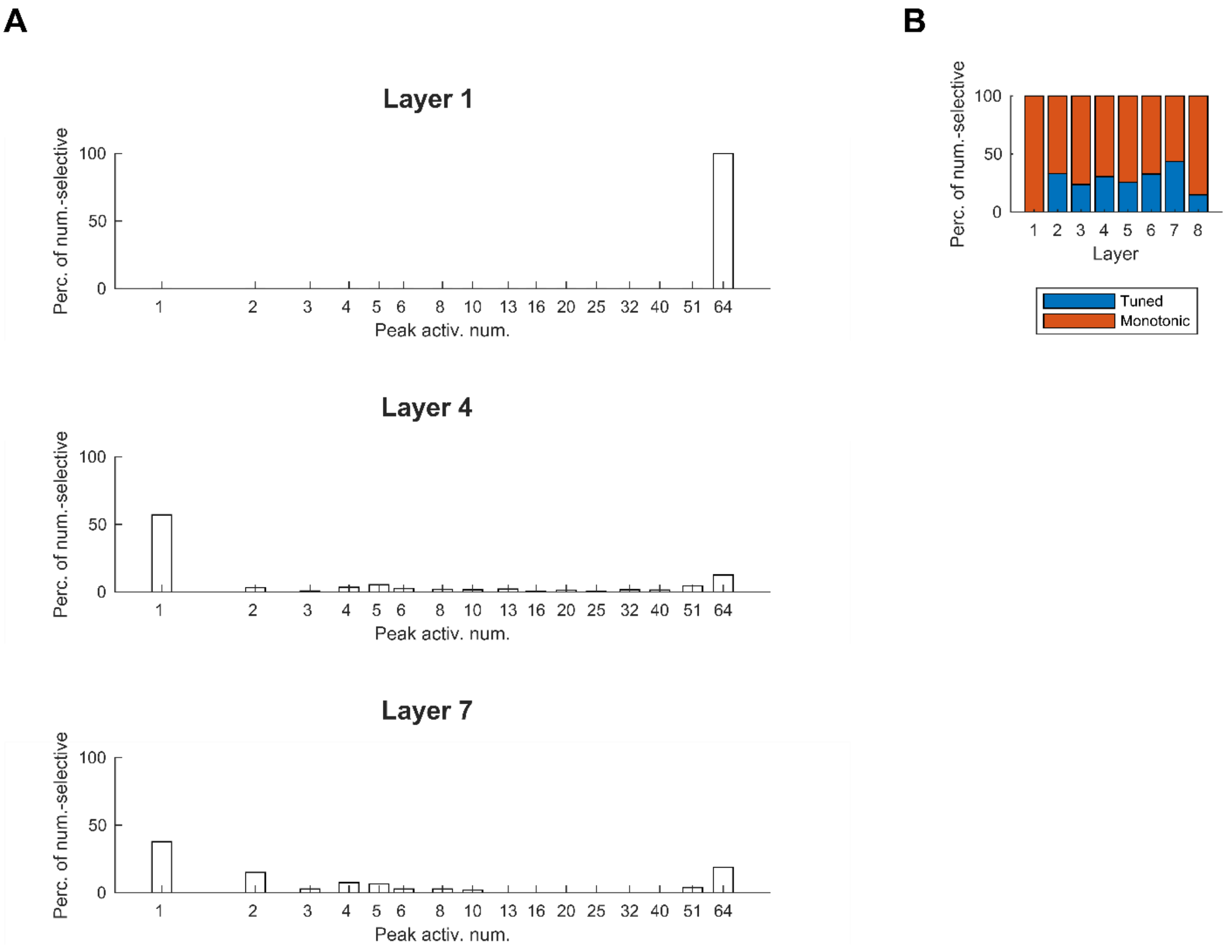


**Figure S3. Most numerosity-selective units are monotonic, but many are tuned.** We tested the numerosity-selective units that responded maximally to four items in the primary analysis (Figure 2A, middle row) on another stimulus set ranging from numerosities 1-5. (A) Histogram plotting the percentage of numerosity-selective units that were maximally activated by each numerosity across both stimulus sets. (B) The percentage of numerosity-selective units that were monotonic (responded maximally to 1 or 64) vs. tuned (responded maximally to an intermediate numerosity) in each layer.

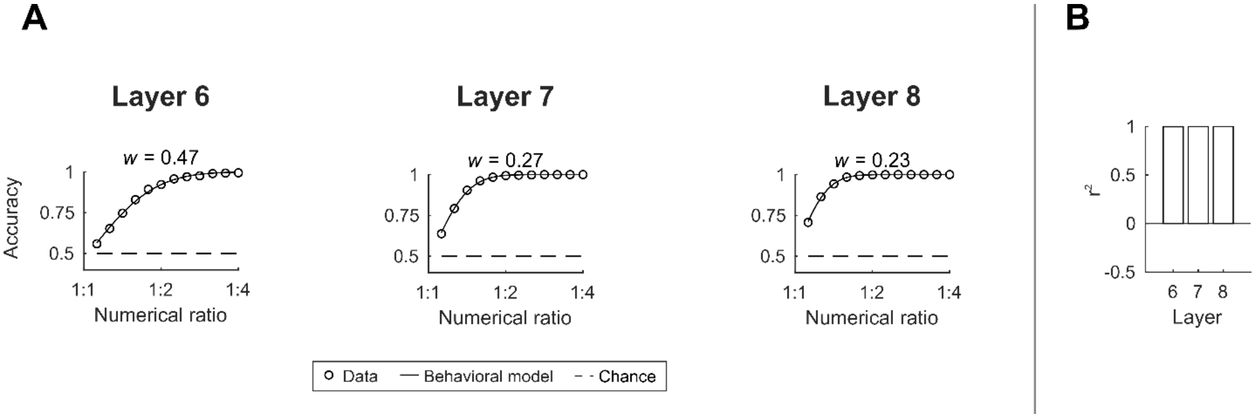


**Figure S4. The empirically validated psychometric function fits the classification accuracy data from the fully connected layers even when more difficult numerical ratios are presented.** (A) Classifier accuracy plotted over numerical ratio for the more difficult stimulus set. Accuracy data is from the three fully connected layers. The line indicates the best-fit psychometric function and *w* indicates the value of the single free parameter. (B) The $r^2$ of the model fit in each layer.

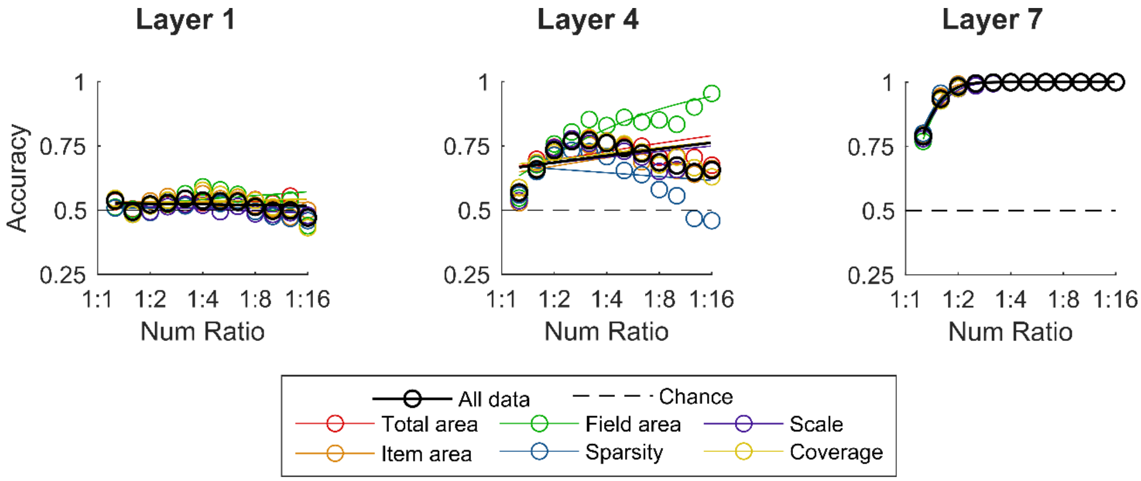


**Figure S5. Numerosity classification performance becomes less affected by non-numerical visual features higher in the DCNN hierarchy.** Circles indicate data and lines are the best fit psychometric curve. Black circles and lines are the same data presented in Figure X for reference. Colored circles and lines correspond to data obtained by testing the classifier on subsets of the validation set in which numerosity varied while the corresponding visual feature was held constant.

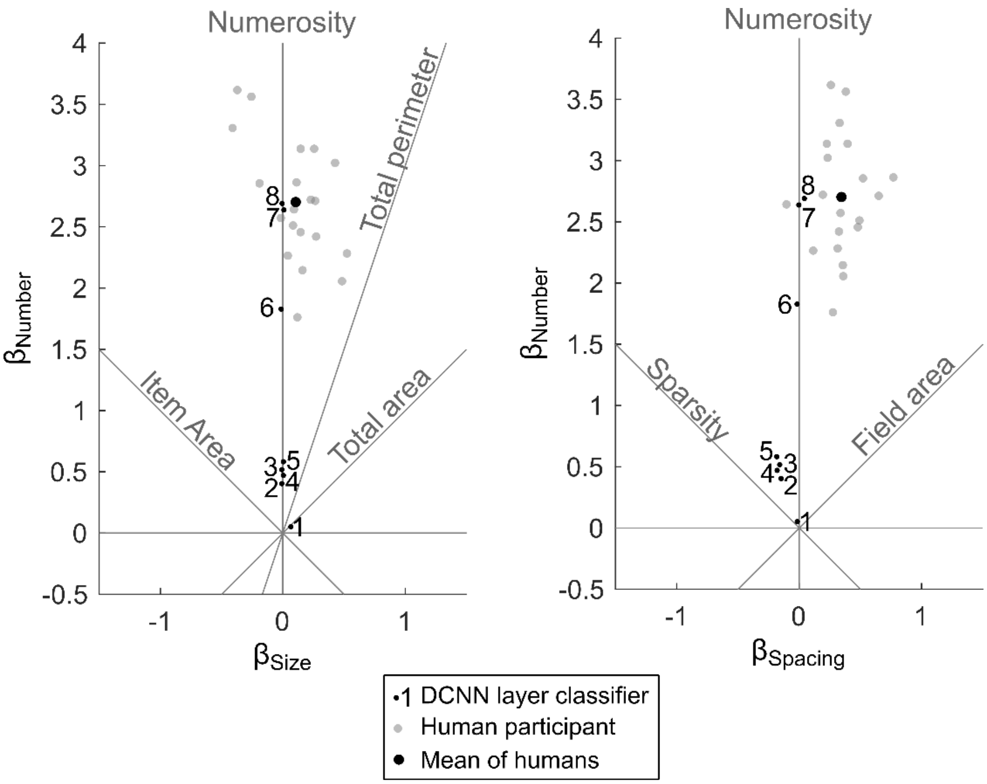


**Figure S6. Layers 7 and 8 perform very similarly to adult humans on numerical ordinal comparison.** Coefficient estimate vectors obtained by fitting the number, size, spacing model to the pair-wise classification data from each layer (DeWind, Adams, Platt, & Brannon, 2015). $\beta_{Num}$ and $\beta_{Size}$ (left panel) and $\beta_{Num}$ and $\beta_{Spacing}$ (right panel) plotted against each other. Gray "feature lines" indicate the expected location of coefficient vectors if that feature were driving classification performance. Unbiased numerical comparison would fall on the center line in both panels (significant positive effect of numerical ratio, but no effect of size or spacing ratio). Layer 1 falls at the origin in both plots indicating that classifications were not driven by any identified feature. The other convolutional layers are largely unaffected by the size of the elements but are biased away from pure numerosity comparison by sparsity. Layer 6 performs unbiased classification of numerosity and has higher acuity (larger $\beta_{Num}$) than the worst human participant (gray points) but is below mean human acuity (larger black point). Layers 7 and 8 perform at human levels of acuity and are similarly unbiased by element size but are less biased by spacing than the average human.

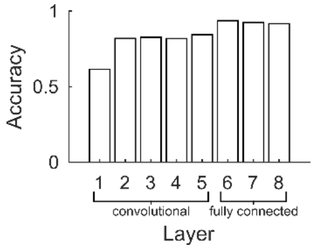


**Figure S7. Classification accuracy on numerosities outside the training set.** A classifier was trained to discriminate which values were greater than 16 among the numerosities 4, 6, 10, 25, 40, and 64 and then tested on numerosities 5, 8, 13, 20, 32, and 51. Mean accuracy on the testing set is plotted for each layer. All p-values << 0.001 (binomial test).

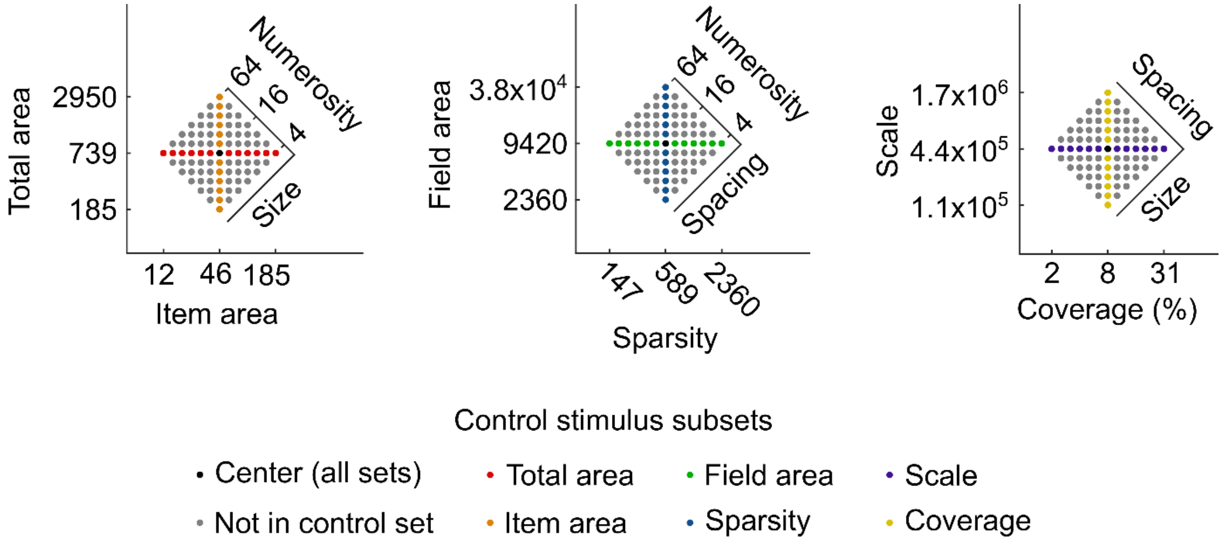


**Figure S8. The 4-64 stimulus set sampling of the stimulus space.** Two-dimensional projections of the full three-dimensional stimulus space. From left to right the collapsed dimension is spacing, size, and numerosity. Each point represents a stimulus. In subsets of the stimuli one visual feature is fixed while numerosity varies across all 13 values. These subsets are color coded by the feature that was fixed: total area (red), item area (orange), field area (green), sparsity (blue), scale (purple), and coverage (yellow). The stimulus at the center of the stimulus space is in all control subsets (black). Many stimuli are not in any subset (gray). Areas (total area, item area, field area, and sparsity) are given in pixels$^2$. Coverage is given as a percentage. Scale is in pixels$^4$; the raw values of scale are difficult to interpret, but scale can be thought of as varying with the “loom” or “zoom” of the stimulus.
